# A chemical method to sequence 5-formylcytosine on RNA

**DOI:** 10.1101/2021.11.23.469780

**Authors:** Ang Li, Xuemeng Sun, A. Emilia Arguello, Ralph E. Kleiner

**Affiliations:** Department of Chemistry, Princeton University, Princeton, NJ 08544

## Abstract

Epitranscriptomic RNA modifications can regulate biological processes, but there remains a major gap in our ability to identify and measure individual modifications at nucleotide resolution. Here we present Mal-Seq, a chemical method to sequence 5-formylcytosine (f^5^C) modifications on RNA based upon selective and efficient malononitrile-mediated labeling of f^5^C residues to generate adducts that are read as C-to-T mutations upon reverse transcription and PCR amplification. We apply Mal-Seq to characterize the prevalence of f^5^C at the wobble position of mt-tRNA(Met) in different organisms and tissue types and find that high-level f^5^C modification is present in mammals but lacking in lower eukaryotes. Our work sheds light on mitochondrial tRNA modifications throughout eukaryotic evolution and provides a general platform for characterizing the f^5^C epitranscriptome.

---

The function of cellular RNA can be modulated by chemical modifications installed post-transcriptionally. Known as the epitranscriptome, over 150 distinct modifications have been reported to exist on RNA^1–2^. A number of well-studied modifications have important roles in RNA metabolism, protein translation, and RNA trafficking^3^, however, we lack information on the function and distribution of most modifications. Further, RNA modification levels and their associated writer, eraser, and reader proteins can be dysregulated in certain disease states^4^, underscoring the need for a comprehensive understanding of epitranscriptomic mechanisms in biological systems.

A major challenge in the study of RNA modifications is the ability to map modifications at single-nucleotide resolution and measure their stoichiometry^5^. While Next-Generation Sequencing (NGS) has revolutionized transcriptomic studies, many modifications are “silent” upon RNA-seq analysis since they are reverse transcribed like the parent unmodified base, necessitating the development of alternative approaches for modification-specific sequencing. Approaches for modification mapping compatible with Illumina sequencing^6^ (the most commonly utilized NGS platform) generally fall into two categories: 1) antibody enrichment of modified RNAs^7–8^ or 2) chemical/enzymatic conversion of the modified base to an adduct that can be identified based upon a distinctive reverse transcription (RT) signature^9–11^. The second approach, particularly when the signature is a sequence mutation as opposed to an RT stop, is often preferred since it can provide higher resolution, less sequence bias, and modification stoichiometry; however, current strategies for RNA modification mapping mediated by chemical or enzymatic conversion are only applicable to a small number of modified bases, and there is a great need for the development of new approaches to characterize the epitranscriptome.

Herein, we develop a chemical approach to sequence 5-formylcytosine (f^5^C) on RNA at single-nucleotide resolution. 5-formylcytosine has been found on isolated tRNA isoacceptors^12–14^, but we lack robust approaches to quantitatively sequence this modification and characterize its distribution across the transcriptome. Our strategy, which we name Mal-Seq, is based upon selective malononitrile-mediated labeling (Fig. 1a) and C-to-T conversion upon reverse transcription, amplification and sequencing. Chemical labeling with malononitrile is mild, efficient, and quantitative, and we exploit these properties to measure the levels of f^5^C at C34 on mt-tRNA(Met) in diverse organisms and tissue types.

**Figure 1.**
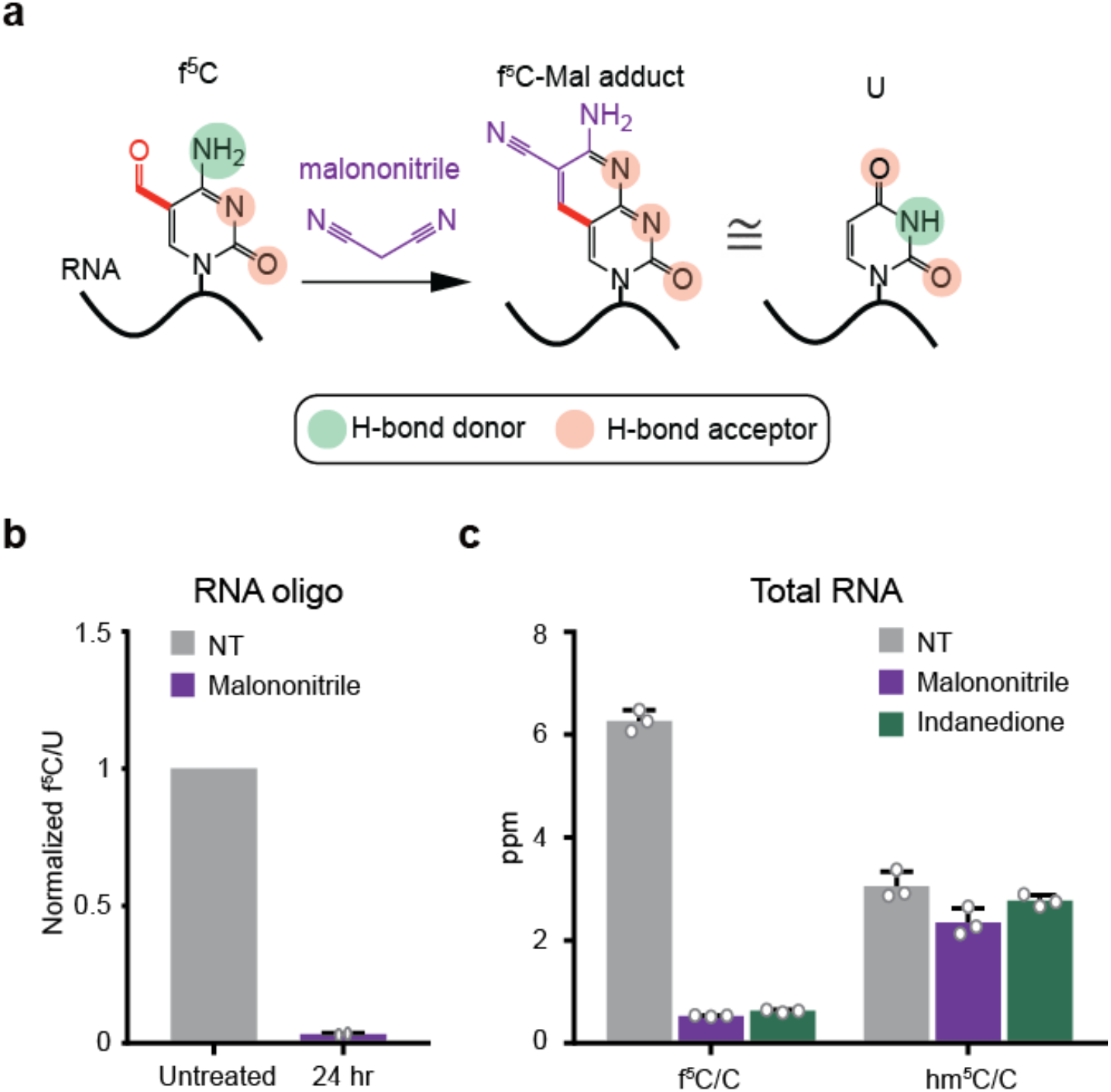
Malononitrile labeling of f^5^C on RNA. (a) Structure of malononitrile-f^5^C adduct and effects on Watson-Crick base pairing. (b) LC-QQQ-MS analysis of f^5^C levels in RNA oligo1 after treatment with malononitrile. Data are mean ± s.d. (*n* = 2). (c) LC-QQQ-MS analysis of f^5^C and hm^5^C levels in total RNA before and after treatment with malononitrile or 1,3-indandione. Data are mean ± s.d. (*n* = 3).

In order to sequence RNA f^5^C, we surveyed the literature for chemical transformations that would be selective for the modified base and generate a mutational signature. Notably, Yi and co-workers previously demonstrated that malononitrile^15^ and 1,3-indandione^16^ react with 5-formylcytosine in DNA to form adducts that induce C-to-T mutations upon DNA polymerase read-through (Fig. 1a and Fig. S1a). Therefore, we investigated the suitability of these reactions for sequencing f^5^C on RNA using total RNA and an artificial f^5^C-containing RNA transcript generated by *in vitro* transcription as model substrates. We started by quantifying depletion of f^5^C in RNA upon treatment with malononitrile or 1,3-indandione using nucleoside LC-MS/MS (Table S1). Gratifyingly, we measured 96.9 ± 0.004% reduction in f^5^C levels upon treatment of our model f^5^C RNA with malononitrile (Fig. 1b). In addition, treatment of total cellular RNA with malononitrile or 1,3-indandione resulted in 90-92% (malononitrile: 91.5 ± 0.002%, 1,3-indandione: 90.0 ± 0.004%) reduction of f^5^C levels (Fig. 1c). Importantly, levels of related modifications in total RNA such as 5-methylcytidine (m^5^C) or 5-hydroxymethylcytidine (hm^5^C) remained unchanged (Fig. 1c and Fig. S1c), and analysis of RNA integrity using gel electrophoresis or Bioanalyzer assay demonstrated minimal RNA degradation (Fig. S1d and S1e), indicating that these transformations are selective for f^5^C and sufficiently mild for RNA sequencing.

Next, we tested whether the generated f^5^C adducts would produce a sequence mutation upon reverse transcription PCR (RT-PCR). Given comparable reaction efficiency between f^5^C and malononitrile or 1,3-indandione, we chose to work with malononitrile due to its enhanced solubility. We treated an *in vitro* transcribed RNA containing a single f^5^C site at 100% stoichiometry with malononitrile and performed RT-PCR. The f^5^C site was positioned in a Taqα1 digestion site such that mutation of C to another base could be monitored by restriction enzyme digestion^17^ (Fig. 2a). As shown, the RT-PCR products generated from untreated f^5^C RNA or an unmodified RNA are quantitatively digested by Taqα1, while malononitrile treatment of the f^5^C RNA inhibited digestion by ~50-60% (Fig. 2b and Fig. S2). To characterize the nature of the sequence change, we performed Sanger sequencing, which indicated that 60% of the transcripts contained a C-to-T mutation, while the remaining 40% contained a C (Fig. 2c). No other mutations were detected at the f^5^C site or surrounding residues. To further confirm our result and generate a calibration curve relating f^5^C stoichiometry and C-to-T conversion, we used high-throughput sequencing. Our data show a linear correlation between f^5^C levels and malononitrile-induced C-to-T mutation with a conversion factor of 0.56 (i.e. 56% C-to-T conversion corresponds to 100% f^5^C) (Fig. 2d and Table S2). Given the depth of coverage afforded by high-throughput sequencing analysis, we could also detect mutations to G/A or deletions at the f^5^C site after malononitrile treatment, but the frequency of such events was low (2.6-3.9%). In addition, we tested an RNA containing two f^5^C modification sites within different sequence contexts and observed similar levels of C-to-T conversion upon malononitrile treatment at each site (Fig. 2e and Table S3). Taken together, our results demonstrate that malononitrile-induced C-to-T mutations can be used to quantitatively sequence f^5^C modifications at nucleotide resolution within RNA. We named this approach Mal-Seq.

**Figure 2.**
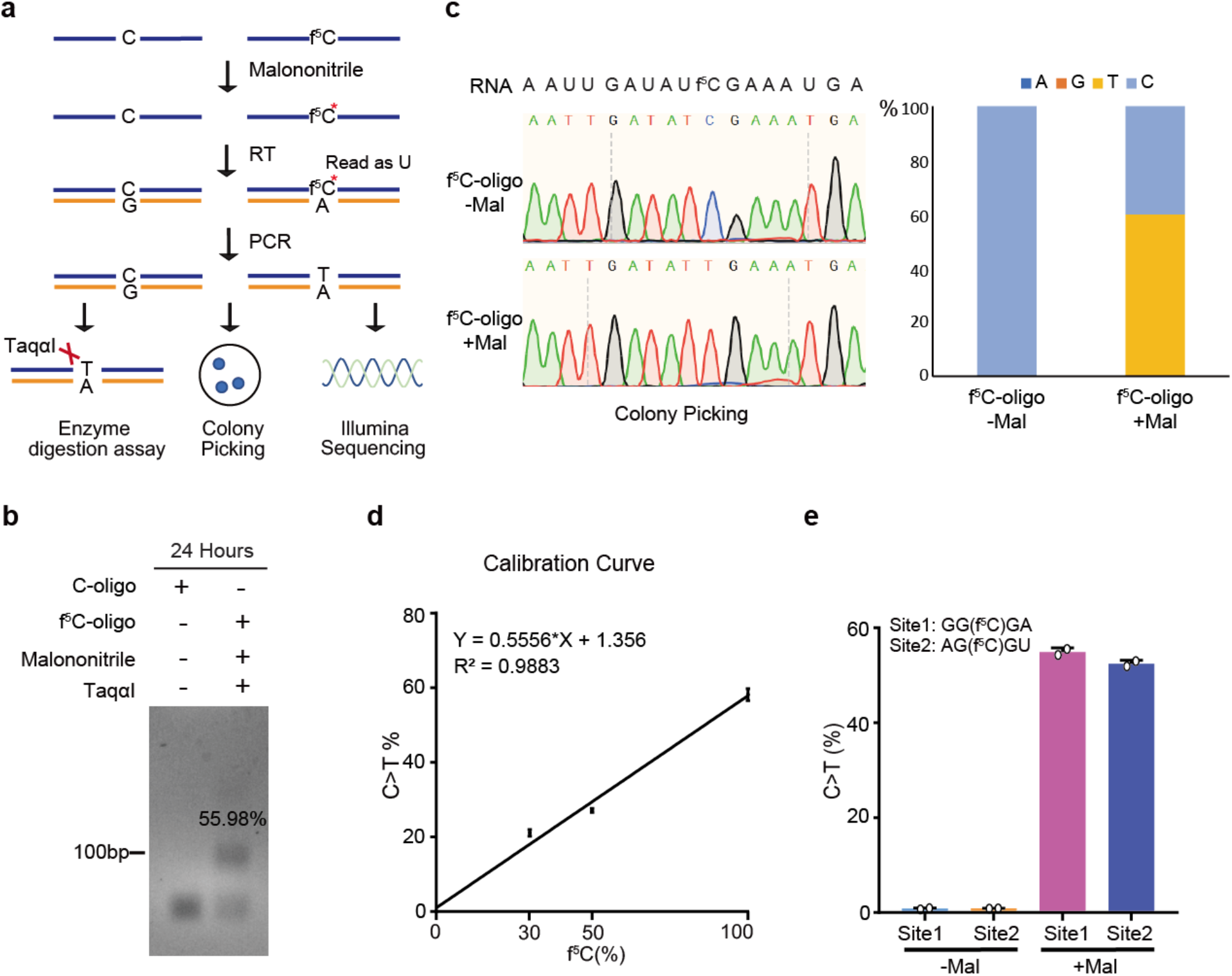
Quantitative sequencing of RNA f^5^C by Mal-Seq. (a) Schematic of Mal-Seq workflow. (b) Taqα1 enzymatic digestion assay to detect base mutations mediated by malononitrile labeling. Malononitrile-mediated C-to-T conversion measured by colony picking and Sanger sequencing. Left: Representative Sanger sequencing traces; Right: Quantification analysis (*n* = 10). (d) Calibration curve relating f^5^C levels in RNA oligo1 and malononitrile-mediated C-to-T conversion. C-to-T mutations were measured by Illumina sequencing. Data are mean ± s.d. (*n* = 2). (e) C-to-T conversion at two distinct f^5^C-modified sites in RNA oligo2. Mutations were measured by Illumina sequencing. Data are mean ± s.d. (*n* = 2).

In order to demonstrate the utility of Mal-Seq, we applied our approach to characterize endogenous f^5^C modification levels in the anticodon loop of mt-tRNA(Met). Studies have indicated the presence of f^5^C at the C34 “wobble base” of mt-tRNA(Met) in a number of organisms, where it is proposed to facilitate the decoding of unconventional AUA and AUU Met codons among mitochondrial genes^18^. However, the lack of a unified, quantitative approach to characterize f^5^C modification levels has led to disparate findings regarding the prevalence and stoichiometry of f^5^C levels in biological systems. We started by applying Mal-Seq to quantify f^5^C on the wobble base of mt-tRNA(Met) from cultured human cells, where this modification has been best studied. Multiple groups have shown that f^5^C biogenesis at this position requires the sequential action of m^5^C methyltransferase NSUN3 followed by Fe(II), α-KG-dependent dioxygenase ALKBH1^12–13, 19–20^, however quantification of modification levels has varied. Suzuki and co-workers used LC/MS analysis to show that C34 is fully modified to f^5^C^20^, while two independent reports relying upon primer extension and bisulfite-based sequencing methods found a mixture of f^5^C and m^5^C at this position^13,19^. Therefore, we extracted total RNA from WT HEK293T cells and performed Mal-Seq using targeted RT-PCR of the anticodon region of mt-tRNA(Met). Our analysis shows 57.98 ± 0.16% malononitrile-induced C-to-T conversion, indicating that mt-tRNA(Met) is fully modified with f^5^C at the wobble base (Fig. 3b, 3c and Table S4), consistent with Suzuki’s findings. In addition, we performed parallel analyses on RNA extracted from ALKBH1 or NSUN3 KO cells generated by CRISPR/Cas9 technology and found <0.3% C-to-T mutation (Fig. 3c, Fig. S3 and Table S4), confirming that both enzymes are required for f^5^C installation on mt-tRNA(Met).

**Figure 3.**
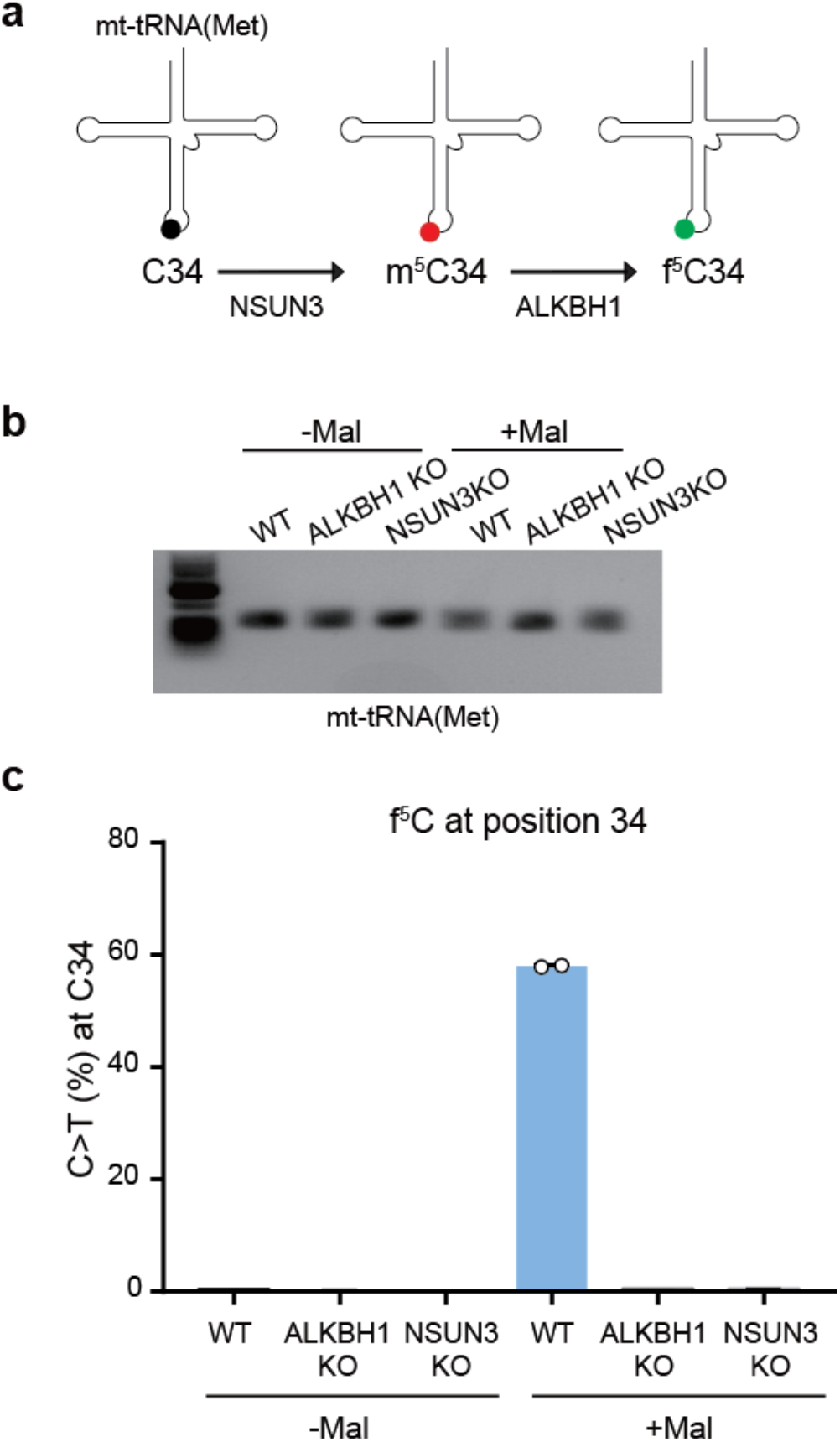
Mal-Seq reveals f^5^C34 in mt-tRNA(Met) is fully modified in HEK293T cells. (a) Schematic showing the formation of f^5^C by NSUN3 and ALKBH1 on mt-tRNA(Met). (b) RT-PCR of mt-tRNA(Met) amplified from WT, ALKBH1 KO, and NSUN3 KO cell lines. (c) C-to-T mutation at C34 on mt-tRNA(Met) detected by Mal-Seq using RNA from WT, ALKBH1 KO, and NSUN3 KO cells. Data are mean ± s.d. (*n* = 2).

We next characterized the presence of f^5^C on the wobble position of mt-tRNA(Met) in other eukaryotes. This modification has been found in organisms including squid^21^, flies^22^, chicken^23^, - and cow^14^, but the extent of f^5^C modification in these species is largely unknown. We obtained total RNA from budding yeast, flies*, C. elegans* and mouse and characterized f^5^C levels by Mal-Seq using species-specific primers for each mt-tRNA(Met) (Fig. S4). In yeast and flies, we found no evidence of f^5^C on mt-tRNA(Met), indicating that this modification is absent or below our limit of detection (Fig. 4a and Table S5). Budding yeast lack a clear ALKBH1 homolog, which is consistent with low f^5^C modification. While f^5^C on mt-tRNA(Met) has been reported to occur in flies, modification levels were partial and quantitation was never performed^22^. In contrast, *C. elegans* showed 27.4 ± 3.3% Mal-Seq C-to-T conversion corresponding to 48.8 ± 5.9% of the mt-tRNA(Met) modified with f^5^C at the wobble position. This is in line with the recent characterization of a mitochondrial ALKBH1 homolog in this organism^24^. We also measured f^5^C levels in different mouse tissues including heart, brain, and liver. In these tissues, we observed 37.8-42.9% (liver: 37.8 ± 1.7%, heart: 41.5 ± 3.2%, brain: 42.9 ± 1.3%) C-to-T conversion corresponding to 67.5-76.6% f^5^C modification, with a slight decrease in the liver (Fig. 4b and Table S5).

**Figure 4.**
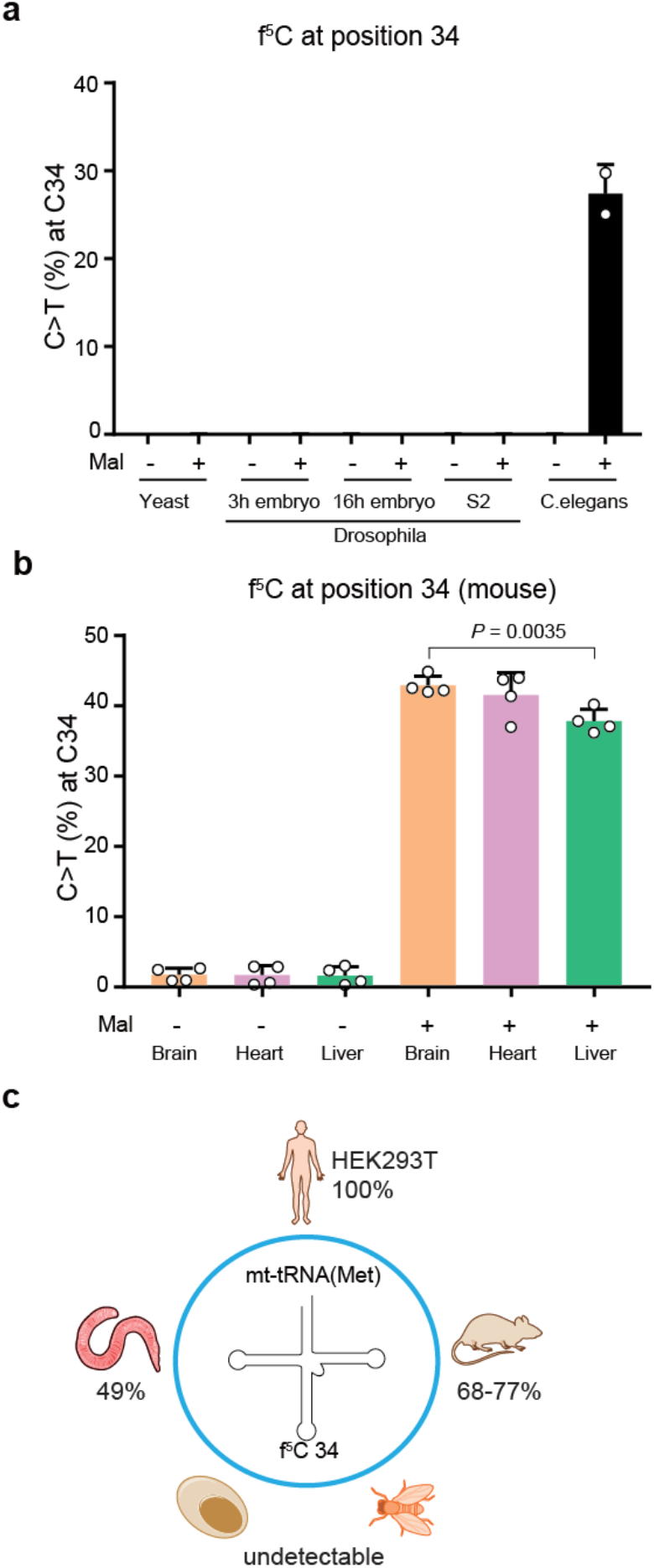
Detection of f^5^C34 on mt-tRNA(Met) among different organisms. (a) Mal-Seq analysis of C34 on mt-tRNA(Met) using RNA extracted from *C.elegans*, budding yeast, and *D. melanogaster*. (b) Mal-Seq analysis of C34 on mt-tRNA(Met) in different mouse tissues. (c) Schematic showing f^5^C levels on mt-tRNA(Met) in different organisms detected by Mal-Seq. Data are mean ± s.d. (*n* = 2 for yeast, *C. elegans*, fly S2 cells; *n* = 4 for mouse tissues; *n* = 2 technical replicates for fly embryos). P values were determined using a two-sided unpaired student’s t-test.

In this work, we develop a chemical sequencing approach, Mal-Seq, for detecting and quantifying 5-formylcytosine on RNA. Mal-Seq analysis of f^5^C modification at the wobble base of mt-tRNA(Met) allows measurement of this modification in different organisms and different tissues types. Our results show that mt-tRNA(Met) is fully modified with f^5^C in human HEK293T cell, and demonstrate on average 72.8 ± 4.7% modification levels in the mouse tissues that we assayed, consistent with the important role of this modification for mitochondrial translation in mammals^12–13, 19^. Interestingly, modification levels are largely invariant in the different murine tissues that we sampled. Oxidation of m^5^C to f^5^C on mt-tRNA(Met) requires ALKBH1, which uses O_2_ and α-KG as cofactors. In principle, ALKBH1 activity (and as a consequence mitochondrial translation efficiency) could be responsive to fluctuations in levels of these central metabolites; while this hypothesis remains to be tested explicitly, our data suggests that installation of f^5^C on mt-tRNA(Met) is largely independent of the physiological fluctuations in metabolite levels across different tissues under the conditions tested. In addition, we find that high-level f^5^C modification (>50%) is characteristic of mammals and absent in lower eukaryotes (Fig. 4c). Since recognition of mitochondrial Met AUA/AUU codons is important in many eukaryotes, other mechanisms must exist to support this role in systems lacking f^5^C. Alternatively, lower level f^5^C modification may be sufficient to satisfy the requirements of mitochondrial translation in these organisms. Finally, the development of a nucleotide resolution sequencing strategy for detecting f^5^C modification opens opportunities for mapping this modified base transcriptome-wide in different organisms. Notably, nucleoside LC-MS analysis has been used to support the existence of f^5^C in yeast mRNA^25^ but individual modification sites have not been reported. In addition, ALKBH1 has been shown to reside outside of the mitochondria in the nucleus^26^, suggesting that f^5^C sites may also exist on non-mitochondrial RNAs in mammals. Our sequencing approach, together with identification of the relevant writer enzymes, should enable comprehensive investigation of the f^5^C epitranscriptome and shed light on its role in biology. Such studies are underway and will be reported in due course.

## Supporting Information

Supplemental figures and methods are available online.

## Acknowledgements

We thank Elizabeth Gavis for providing total RNA from fly tissues, Zihong Chen and Joshua Rabinowitz for providing mice tissues, Anuj Sharma and Andrew Leifer for providing worms, and Eric Lai for providing fly S2 cells. This research was supported by the National Institutes of Health (RO1 GM132189 to R.E.K.), the National Science Foundation (CAREER award MCB-1942565 to R.E.K.), Sidney Kimmel Foundation, and Alfred P. Sloan Foundation. A.E.A. acknowledges support from an Eli Lilly-Edward C. Taylor Fellowship in Chemistry and A.L. was supported by the Princeton Catalysis Initiative. All authors thank Princeton University for financial support.

