## Supplemental Information for "A chemical method to sequence 5-formylcytosine on RNA"

Table of contents

i. Methods
ii. Supplementary Figure 1
iii. Supplementary Figure 2
iv. Supplementary Figure 3
v. Supplementary Figure 4
vi. Supplementary Figure 5
vii. Supplementary Table1: Dynamic multiple reaction monitoring (DMRM) parameters of for quantification nucleosides by triple quadrupole (QQQ) LC-MS/MS.
viii. Supplementary Table 2: Reads information of Mal-seq calibration curve.
ix. Supplementary Table 3: Reads information of Mal-seq using RNA oligo2.
x. Supplementary Table 4: Mt-tRNA(met) reads information of Mal-seq using RNA from WT, ALKBH1 and NSUN3 knockout HEK cells.
xi. Supplementary Table 5: Mt-tRNA(met) reads information of Mal-seq using RNA from Yeast, C.elegans and Fly tissues.
xii. Supplementary Table 6: Oligonucleotides and primers.

### Methods

*In vitro transcription*

Model RNA oligos containing C or $f^5C$ were generated using T7 RNA polymerase in 100 µL reaction mix containing 50 mM Tris-HCl pH 7.5,15 mM $MgCl_2$, 5 mM dithiothreitol (DTT), 2 mM spermidine, 1 µL RNase Inhibitor (NEB),1 mM ATP, UTP, GTP, and 1 mM CTP or 5-formylcytidine-5'-triphosphate (Trilink Biotech). After 2 hr incubation at 37°C, 10 U of RNase-free DNase I was added and the reaction was incubated at 37 °C for 30 min. 5 µL of 500 mM EDTA was added to stop the reaction. RNA was purified by phenol-chloroform extraction and ethanol precipitation.

*Malononitrile sequencing*

500 ng of RNA oligo or 3 µg total RNA was incubated with 150 mM of malononitrile in 30µL 1X TE (pH = 7.4) buffer containing 1 µL RNase Inhibitor at 37°C for 24 hr. After reaction, 5 µL reaction mix were incubated with 1 µL 10 µM RT primer and 1 µL 10 mM ATP at 65°C for 5 min.The mix then used for reverse transcription with Superscript II following the manufacturer's instructions. cDNA was then PCR amplified by AccuPrime™ Pfx DNA Polymerase (Thermo) and PCR products were gel-purified and submitted for NGS (Genewiz Amplicon Sequencing) or digested with restriction enzymes for cloning, colony picking, and Sanger sequencing.

*Enzyme digestion assay*

PCR products were incubated with 4 U of Taq$\alpha$1 restriction enzyme (NEB) in 1X CutSmart buffer (NEB) at 65 °C for 1 hr and analyzed by 2% agarose gel electrophoresis.

*Nucleoside LC-MS*

In general, after malononitrile reaction, total RNA was purified using Zymo RNA clean kit. RNA (1-3 µg) was then digested in 20 µL with 2 U of Nuclease P1 (Wako) and final concentrations of 7 mM NaOAc and 0.4 mM $ZnCl_2$ at 37 °C for 2 hr. The digested mixture was subjected to dephosphorylation in a total volume of 30 µL with 2 U of Antarctic Phosphatase (NEB) and a final concentration of 1X Antarctic Phosphatase buffer at 37 °C for 2hr. Quantification of ribonucleosides from biological samples by triple quadrupole (QQQ) liquid chromatography-tandem mass spectrometry (LC-MS/MS) was performed on an Agilent 1260 LC Infinity II system coupled to an Agilent 6470 LC/TQ module on a Hypersil GOLD aQ column (Thermo, #25303-152130). Solvent flow rate was kept at 0.4 mL/min, and the pressure limit was set at 600 bar. LC-MS-grade acetonitrile (Fisher #85188) and (0.1% formic acid (Fisher #A117-50) in ultrapure water were used as mobile phases. The ionization source was set to the following parameters: 350 °C gas temperature, 12 L/min gas flow, 20 psi nebulizer pressure, and 2500 V capillary voltage. Ribonucleoside standards were synthesized or obtained commercially according to Table S1. Analyte detection was done by dynamic multiple reaction monitoring (DMRM) according to the parameters in Supplementary Table 1. Abundant nucleosides in biological samples (ACGU) were quantified in 1 ng of sample, while less abundant ones ($m^5C$, $hm^5C$, $f^5C$, $i^6A$) were analyzed in 100-200 ng of sample. Analytes were separated on a 7 min isocratic flow of 100% 0.1% formic acid, followed by a 3 min post-run unless otherwise noted. Measurement was taken from 3 distinct biological replicates.

*Mutation detection*
Barcode splitter was used to demultiplex the reads. We adapted Breseq[1] Consensus mode for mutation detection (Galaxy Version 0.34.0+1). Briefly, reads were mapped to specific reference sequences by Bowtie2 (2.3.5). The mutation ratio was then calculated by using the following formula: number of mutated reads / total reads.

*Total RNA extraction from mouse tissues*
Mouse studies followed protocols approved by the Princeton University Animal Care and Use Committee. The C57BL/6 mice (10-24 weeks old) were a generous gift from Prof. Joshua Rabinowitz and were housed on a normal light cycle (8AM-8PM) at Princeton University. Tissue harvest was performed after euthanasia by cervical dislocation. Tissues were quickly dissected, clamped with a pre-cooled Wollenberger clamp, and dropped in liquid nitrogen. Frozen tissue was ground by a Cyromill at cryogenic temperature (Retsch, Newtown, PA). Ground tissue was then weighed (~20 mg) for total RNA extraction by adding 200 µL TRIzol.

*Total RNA extraction from Yeast and C. elegans*
*S. cerevisiae* (strain BY4743) were grown in YPD medium and harvested when the OD reached 0.6. *C. elegans* was a generous gift from Prof. Andrew Leifer. Yeast cells and *C. elegans* were homogenized using beads with 1 mL Trizol followed by freeze and thaw (3x).

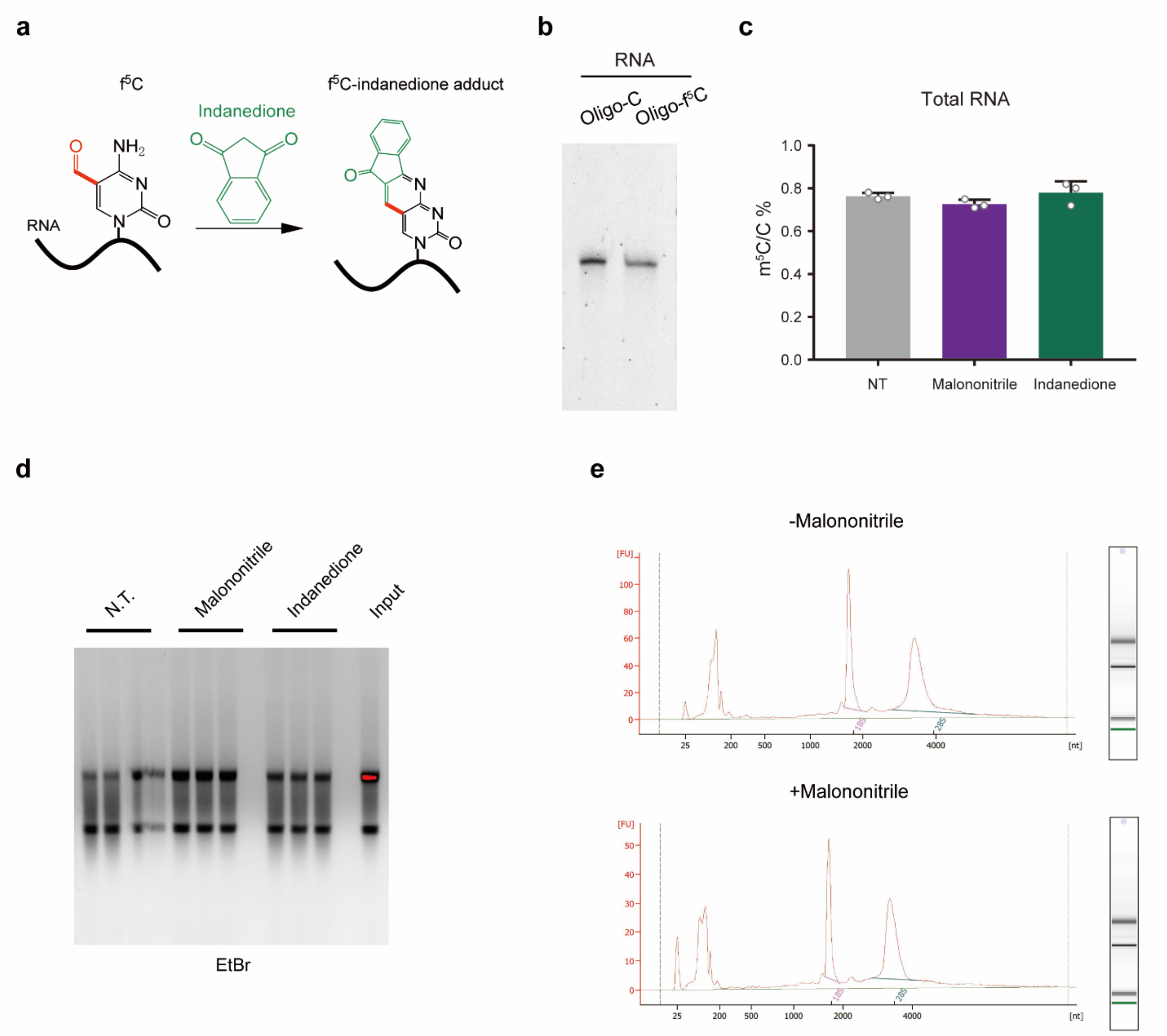


**Figure S1**. (a) Scheme for 1,3-indanedione-mediated labeling of RNA $f^5C$. (b) PAGE gel showing the purity of the IVT constructs. (c) Measurement of $m^5C$ levels in total RNA by LC-QQQ-MS after malononitrile or 1,3-indanedione treatment. Data are the mean ± s.d. (n = 3). (d) Agarose gel showing the integrity of total RNA after malononitrile or 1,3-indanedione treatment. (e) Bioanalyzer results showing RNA integrity after malononitrile treatment.

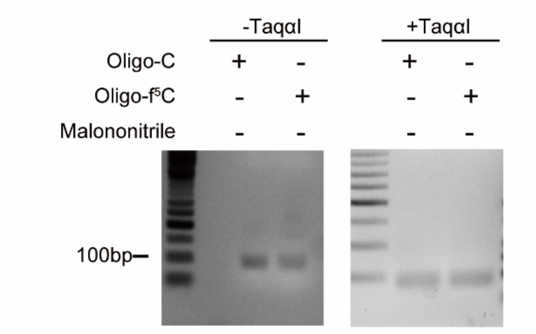


**Figure S2**. Enzyme digestion assay to detect the base change efficiency mediated by malononitrile.

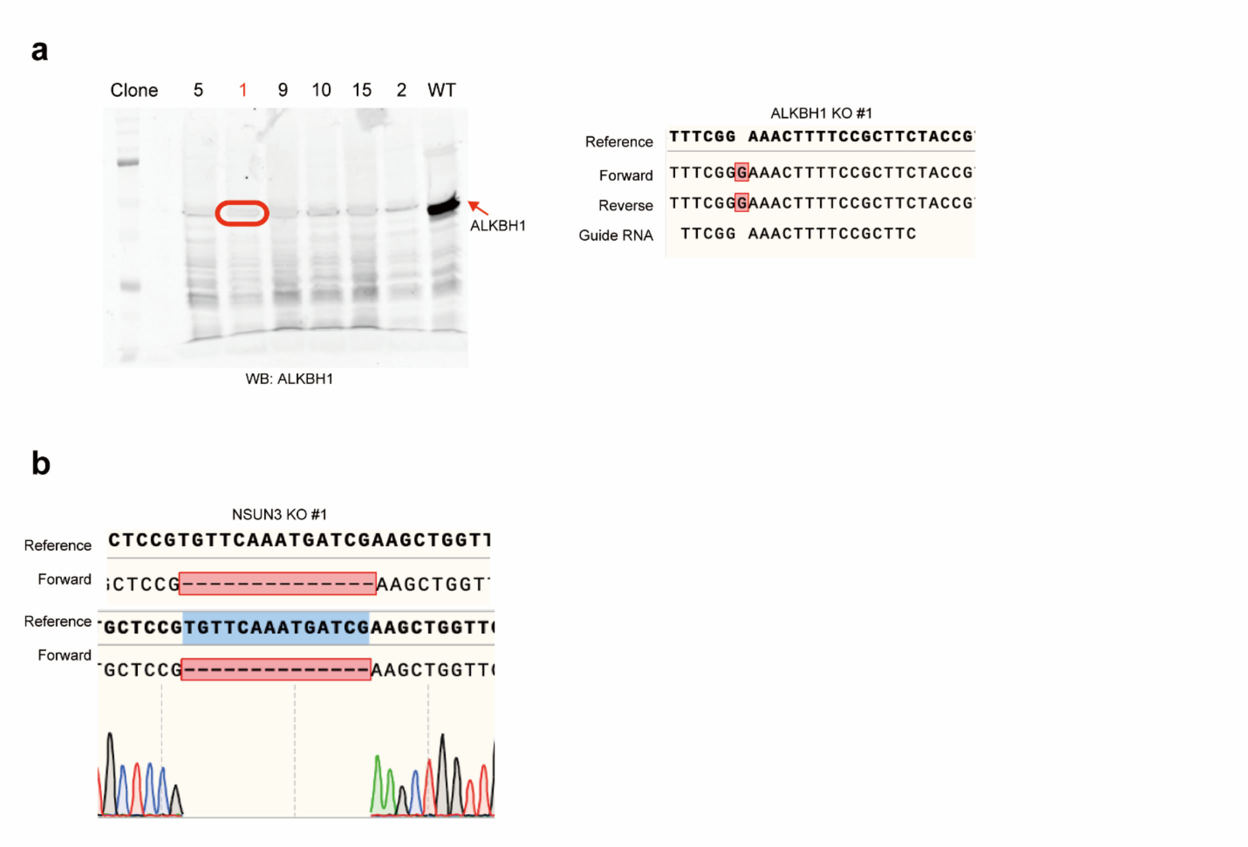


**Figure S3.** Validation of ALKBH1 and NSUN3 KO cells. (a) Western blot (left) and Sanger sequencing (right) showing the ALKBH1 KO. (b) Sanger sequencing showing NSUN3 KO.

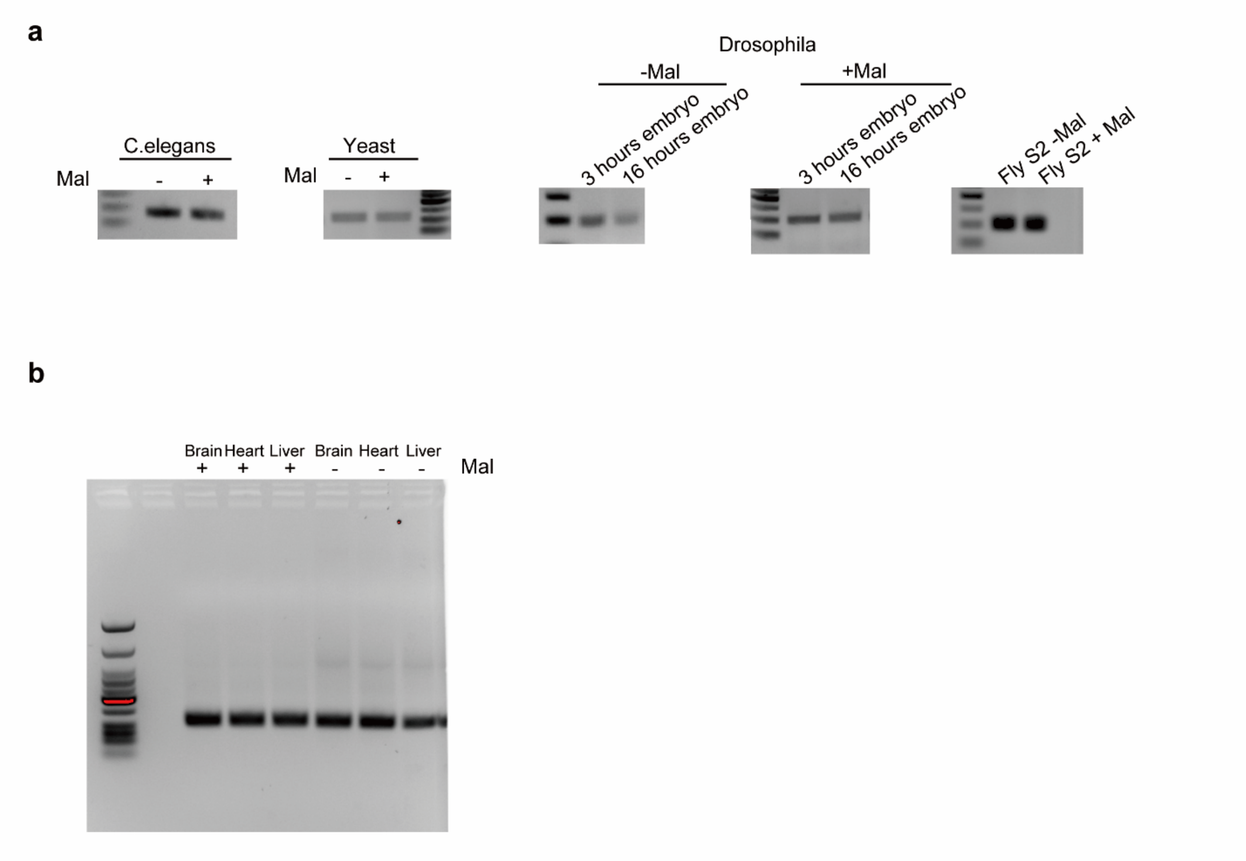


**Figure S4.** (a-b) Agarose gels showing RT-PCR products for mt-tRNA(Met) from RNA isolated from *C.elegans*, yeast, drosophila (a) and murine tissues (b).

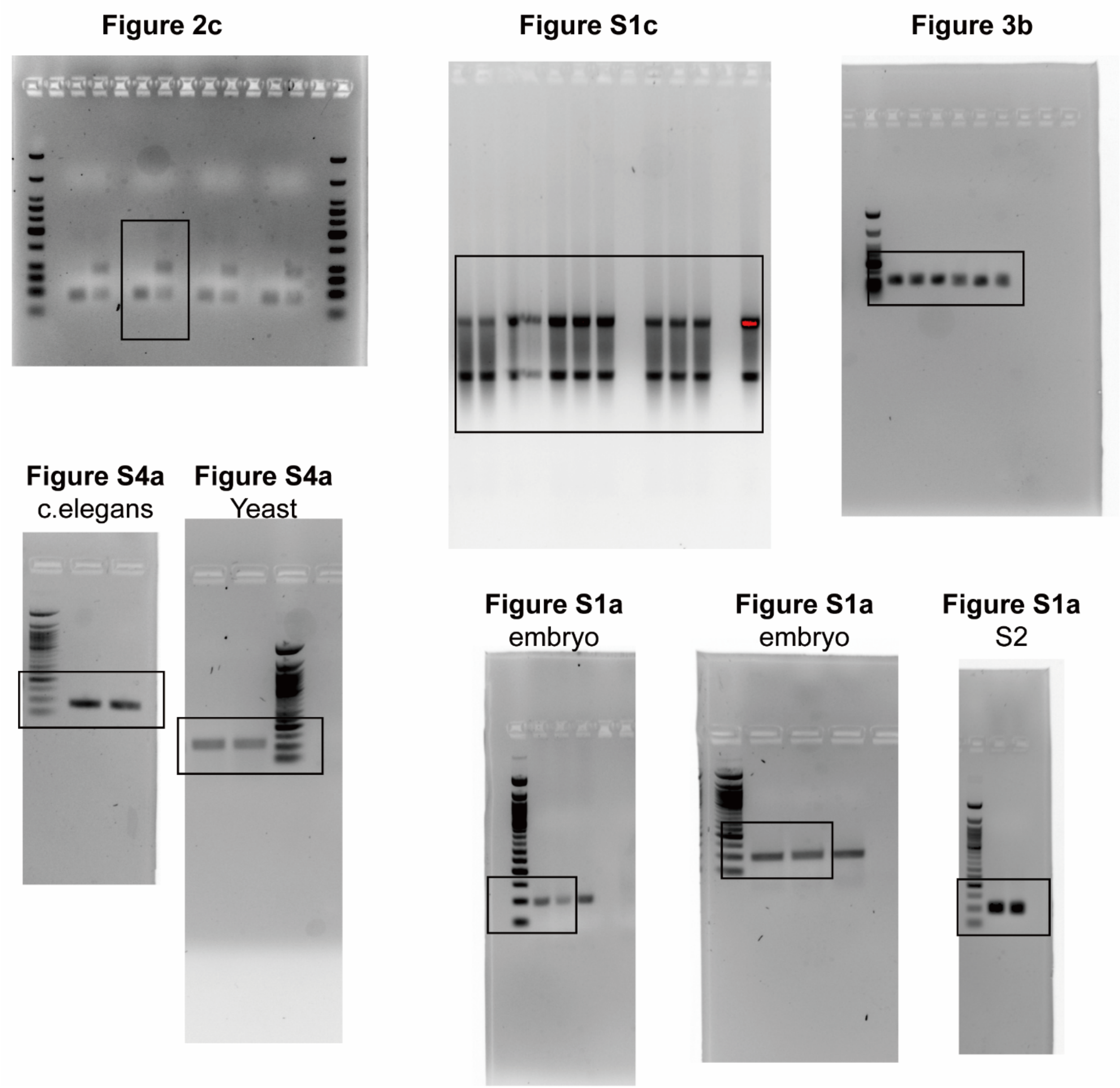


**Figure S5.** Uncropped gel images.

**Table S1**: Dynamic multiple reaction monitoring (DMRM) parameters of for quantification nucleosides by triple quadrupole (QQQ) LC-MS/MS.

| Nucleoside | Fragmentor Energy (V) | Collision Energy (V) | Parent Ion $[M+H]^+$ | Product Ion | Source/Vendor |
|---|---|---|---|---|---|
| A | 100 | 20 | 268.1 | 136.1 | Sigma |
| G | 80 | 13 | 284.1 | 152.1 | Sigma |
| C | 70 | 14 | 244.1 | 112.1 | Sigma |
| $m^5C$ | 70 | 14 | 258.1 | 126.1 | Carbosynth |
| $hm^5C$ | 70 | 14 | 274.0 | 142.0 | Carbosynth |
| $f^5C$ | 70 | 14 | 268.1 | 136.1 | Berry & Associates |
| U | 70 | 7 | 245.1 | 113.1 | Sigma |

**Table S2.** Read information for Mal-Seq calibration curve.

| Samples | Total reads | C | T | C>T(%) |
|---|---|---|---|---|
| oligoC_1 | 710 | 705 | 2 | 0.28169 |
| oligoC_2 | 1254 | 1248 | 0 | 0 |
| 30% oligof5C_1 | 1224 | 929 | 255 | 20.83333 |
| 30% oligof5C_2 | 1074 | 799 | 238 | 22.16015 |
| 50% oligof5C_1 | 729 | 501 | 197 | 27.02332 |
| 50% oligof5C_2 | 568 | 397 | 150 | 26.40845 |
| 100% oligof5C_1 | 1672 | 653 | 963 | 57.59569 |
| 100% oligof5C_2 | 7068 | 2929 | 3999 | 56.57895 |

**Table S3.** Read information for Mal-Seq analysis using RNA oligo2.

| Samples | Total reads | C | T | C>T(%) |
|---|---|---|---|---|
| Oligo2_rep1_f5c1_mal | 3089 | 1317 | 1676 | 54.25704 |
| Oligo2_rep1_f5c2_mal | 3069 | 1331 | 1592 | 51.87357 |
| Oligo2_rep2_f5c1_mal | 3013 | 1244 | 1673 | 55.52605 |
| Oligo2_rep2_f5c2_mal | 2991 | 1316 | 1584 | 52.95888 |
| Oligo2_rep1_f5c1_no-mal | 13111 | 12749 | 90 | 0.686446 |
| Oligo2_rep1_f5c2_no-mal | 13030 | 12587 | 109 | 0.836531 |
| Oligo2_rep2_f5c1_no-mal | 12431 | 12082 | 115 | 0.925107 |
| Oligo2_rep2_f5c2_no-mal | 12337 | 11961 | 110 | 0.891627 |

**Table S4.** Mt-tRNA(Met) read information for Mal-Seq analysis of WT HEK293T, ALKBH1 KO, and NSUN3 KO cells.

| **Samples** | **Total reads** | **C** | **T** | **C>T(%)** |
|---|---|---|---|---|
| WT1_Mal | 22043 | 9113 | 12755 | 57.86417 |
| WT2_Mal | 24499 | 9881 | 14231 | 58.08809 |
| WT1_no-mal | 19176 | 19127 | 28 | 0.146016 |
| WT2_no-Mal | 17109 | 17066 | 8 | 0.046759 |
| ALKBH1_KO1_Mal | 20572 | 20484 | 38 | 0.184717 |
| ALKBH1_KO2_Mal | 838 | 837 | 1 | 0.119332 |
| ALKBH1_KO1_no-mal | 23974 | 23926 | 15 | 0.062568 |
| ALKBH1_KO2_no-Mal | 1431 | 1426 | 1 | 0.069881 |
| NSUN3_KO1_Mal | 3093 | 3090 | 2 | 0.064662 |
| NSUN3_KO2_Mal | 2338 | 2328 | 7 | 0.299401 |
| NSUN3_KO1_no-mal | 3025 | 3023 | 0 | 0 |
| NSUN3_KO2_no-Mal | 2656 | 2656 | 0 | 0 |

**Table S5.** Mt-tRNA(Met) read information for Mal-Seq analysis of RNA from yeast, *C. elegans*, fly tissues, and murine tissues.

| Samples | Total reads | C | T | C>T(%) |
|---|---|---|---|---|
| Yeast_mal1 | 30492 | 30463 | 2 | 0.006559 |
| Yeast_mal2 | 31318 | 31270 | 16 | 0.051089 |
| Yeast no mal1 | 13296 | 13289 | 4 | 0.030084 |
| Yeast no mal2 | 36513 | 36466 | 10 | 0.027388 |
| C.elegans_no_mal1 | 6559 | 6557 | 1 | 0.015246 |
| C.elegans+mal1 | 2159 | 1515 | 642 | 29.73599 |
| C.elegans_no_mal2 | 5937 | 5928 | 2 | 0.033687 |
| C.elegans+mal2 | 890 | 667 | 223 | 25.05618 |
| Flys2-no-mal2 | 2304 | 2301 | 1 | 0.043403 |
| Flys2+mal2 | 36957 | 36944 | 7 | 0.018941 |
| Flys2-no-mal1 | 9914 | 9908 | 2 | 0.020173 |
| Flys2+mal1 | 12666 | 12659 | 6 | 0.047371 |
| fly embyro3 +mal1 | 37046 | 36983 | 4 | 0.010797 |
| fly embyro16 +mal1 | 58778 | 58711 | 16 | 0.027221 |
| fly embyro3-no-mal1 | 10003 | 9995 | 1 | 0.009997 |
| fly embyro16-no-mal1 | 9752 | 9747 | 2 | 0.020509 |
| fly embyro3 +mal2 | 42066 | 42041 | 13 | 0.030904 |
| fly embyro16 +mal2 | 10471 | 10466 | 1 | 0.00955 |
| fly embyro3-no-mal1 | 37150 | 37138 | 4 | 0.010767 |
| fly embyro16-no-mal1 | 2304 | 2301 | 1 | 0.043403 |
| Brain-1+mal | 2320 | 1270 | 1041 | 44.87069 |
| heart-1+mal | 2399 | 1350 | 1049 | 43.72655 |
| liver-1+mal | 503 | 319 | 182 | 36.1829 |
| Brain-2+mal | 2284 | 1316 | 964 | 42.20665 |
| heart-2+mal | 4203 | 2345 | 1848 | 43.96859 |
| liver-2+mal | 4015 | 2397 | 1614 | 40.19925 |
| Brain-3+mal | 3985 | 2276 | 1694 | 42.50941 |
| heart-3+mal | 3836 | 2413 | 1419 | 36.99166 |
| liver-3+mal | 1749 | 1094 | 648 | 37.04974 |
| Brain-4+mal | 3367 | 1945 | 1414 | 41.99584 |
| heart-4+mal | 5624 | 3284 | 2325 | 41.34068 |
| liver-4+mal | 6278 | 3889 | 2374 | 37.81459 |
| Brain-1-no mal | 5770 | 5608 | 153 | 2.651646 |
| heart-1-mo mal | 15825 | 15336 | 461 | 2.913112 |
| liver-1-no mal | 18100 | 17518 | 544 | 3.005525 |
| Brain-2-no mal | 3594 | 3502 | 88 | 2.448525 |
| heart-2-no mal | 8887 | 8617 | 255 | 2.86936 |
| liver-2-no mal | 9113 | 8888 | 214 | 2.348294 |

| | | | | |
|---|---|---|---|---|
| Brain-3-no mal | 3769 | 3727 | 35 | 0.928628 |
| heart-3-no mal | 3275 | 3261 | 12 | 0.366412 |
| liver-3-no mal | 1357 | 1351 | 4 | 0.294768 |
| Brain-4-no mal | 2894 | 2862 | 26 | 0.898411 |
| heart-4-no mal | 5801 | 5761 | 34 | 0.586106 |
| liver-4-no mal | 4926 | 4878 | 40 | 0.812018 |

**Table S6.** Oligonucleotides and primers used in this work.

| Name | Sequence |
|---|---|
| RNA oligo1_template | TTCCCTTACCTACCACTTCCATCACATACTCATTTCGATATCAATT CTATACATACTCTCACTCTCACCTCCCTATAGTGAGTCGTATTA |
| RNA oligo1 sequence | GAGGUGAGAGUGAGAGUAUGUAUAGAAUUGAUAUCGAAAUGAG UAUGUGAUGGAAGUGGUAGGUAAGGGAA |
| RNA oligo2_template | TACTCTATCCTTAACTACATCTACTCATCACGCTAATACTCTATCA TCTCCACATTCCAATATTTCCTAACTATACACCCAAACTTAAAACA TATTCTCAAACCCACCTTCGCCAAACAATCCATACCTCTTAATACT CATCCCTATAGTGAGTCGTATTA |
| RNA oligo2 sequence | GAUGAGUAUUAAGAGGUAUGGAUUGUUUGGCGAAGGUGGGUU UGAGAAUAUGUUUUAAGUUUGGGUGUAUAGUUAGGAAAUAUUG GAAUGUGGAGAUGAUAGAGUAUUAGCGUGAUGAGUAGAUGUA GUUAAGGAUAGAGUA |
| oligo1 Reverse | GGAATTCCTTCCCTTACCTACCACTT |
| oligo1 Forward | CGGATCCGGGAGGTGAGAGTGAGA |
| oligo2 Reverse | TACTCTATCCTTAACTACAT |
| oligo2 Forward | CGGATCCAGTCAAGATGAGTATTAAGAGG |
| human-mt-tRNA-met RT | TAGTACGGGAAGGGTATAACCAACATTT |
| human-mt-tRNA-met forward | AGTAAGGTCAGCTAAATAAGC |
| human-mt-tRNA-met reverse | CGCGGATCCGTACTGACCGGATCCTAGTACGGGAAGGGTATAACCAA C |
| mouse-mt-tRNA-met RT | TAGTACGGGAAGGATTTAAACCAACGTT |
| mouse-mt-tRNA-met forward | CGCGGATCCGAGATTCCAGTAAGGTCAGCTAATTAAG |
| mouse-mt-tRNA-met reverse | ATCACGCCGGAATTCTAGTACGGGAAGGATTTAAA |
| Fly-mt-tRNA-met RT | TAAAAAGAAAAGGATTATAACCTTTATA |
| Fly-mt-tRNA-met forward | CGCGGATCCATTACTCGAAAAAGATAAGCTAATTAAGCT |
| Fly-mt-tRNA-met reverse | CGATGTCCGGAATTCTAAAAAGAAAAGGATTATAACC |
| Yeast-mt-tRNA-met RT | TACTTGTAGAAGGAATTGAACCTTACA |
| Yeast-mt-tRNA-met forward | CGCGGATCCTCCGGAGAGGAATTCGAGCTTGTATAGTTTAA |
| Yeast-mt-tRNA-met reverse | TGACCACCGGAATTCTACTTGTAGAAGGAATTGAA |
| worm-mt-tRNA-met RT | AATAAGAGAAAACCACCTCAAG |
| worm-mt-tRNA-met forward | CGCGGATCCTCCGCGAAAGTAAGATAGGATAATTAAG |
| worm-mt-tRNA-met reverse | CTTGTACCGGAATTCAATAAGAGAAAACCACCT |
